# A genomic test of subspecies in the *Eunota togata* species group (Coleoptera: Cicindelidae): Morphology masks evolutionary relationships and taxonomy

**DOI:** 10.1101/2023.01.05.522877

**Authors:** Robert A. Laroche, Daniel P. Duran, Cin-Ty A. Lee, William Godwin, Stephen J. Roman, David P. Herrmann, Scott P. Egan

## Abstract

Most of the world’s biodiversity is described primarily or exclusively using morphological traits that may not always reflect the true evolutionary units. Accurate taxonomy is critical for conservation efforts and re-evaluation of traditional taxonomy may often be warranted since species and subspecies are frequently the focus of conservation and faunistic studies. Here, we test comprehensive taxonomic hypotheses of morphologically defined subspecies in the tiger beetle, *Eunota togata* (LaFerté-Sénectère, 1841). The four recognized subspecies were delineated based mainly on the dorsal coloration and extent of white markings termed maculations. We combine inferences from mtDNA genealogies and genome-wide multilocus data to elucidate the evolutionary relationships within the group and assess the taxonomic implications. Three of the four subspecific taxa delineated by morphology were not supported by the genomic or mtDNA data. In fact, the species-level diversity in this group was underestimated, as *E. togata* was found to represent three well-supported distinct species in all genetic analyses. Emerging from these analyses, we also document an intriguing example of convergent evolution in lighter colored *E. togata* adapting to similar white sand backgrounds. Our collective work underscores the importance of integrating molecular methods with morphology for species and subspecies delimitation and conservation.

## Introduction

Species delineation has been traditionally based on morphological characters for the vast majority of eukaryotic taxa, with a lesser reliance on behavioral, ecological or other characters (Dayrat 2005). This model is implicitly based on the idea that fixed morphological differences in two or more sets of populations are the result of the splitting of gene pools from a single ancestral taxon. This method of recognizing species as entities that are consistently distinct with respect to body structures is known as the Morphological Species Concept (MSC; Cronquist 1978) and its derivatives (Coyne and Orr 2004). In recent decades, taxonomists have incorporated molecular data into taxonomic revisions and, to a lesser degree, species descriptions (Brandão-Dias et al. 2022), resulting in significant changes to established taxonomic frameworks that are incongruous with the MSC (e.g., Barrowclough et al 2016, Ruiz-Garcia et al 2018, Duran et al. 2019, Titus et al 2019, Kim et al. 2022).

Taxonomic accuracy is essential for effective conservation efforts (Mace 2004). These efforts are typically applied to species and, to a lesser extent, subspecies, which are often recognized as distinct evolutionary lineages worthy of protection (United States 1983, Phillimore and Owens 2006, Haig et al. 2006, Knisley et al. 2014). Hence, the establishment of taxonomy that appropriately represents the divergence between populations is a critical prerequisite to any conservation assessment (Ghisbain et al. 2021). In some cases, taxonomic confusion has led to species being mistakenly included or excluded from conservation efforts, or even erroneously considered threatened or extinct and thus diverting limited resources away from other threatened taxa (May 1990, Russell & Craig 2013, Ely et al. 2017). Further, concerns have been raised that the means by which taxonomic units are defined could potentially influence conservation policy and may lead to inconsistent conservation efforts between biological classes (Garnett & Christidis 2017). Thus, integrating multiple forms of data in taxonomic descriptions is also necessary to provide a foundation for targeted conservation approaches.

Tiger beetles (Cicindelidae) are a charismatic and popular group of insects (Knisley & Schultz 1997, New 2007) with approximately 3,000 species described worldwide (Wiesner 2021). However, the taxonomy of the group has been critically outdated and recent studies have shown that the morphology-based higher-level taxonomy was incongruous with integrative approaches that incorporated both phylogenomic data and morphology (Duran & Gough 2020). Studies of the North American fauna demonstrated that the genus level taxonomy was similarly outdated (Gough et al. 2019; Duran & Gough 2019). Species delineation in tiger beetles has traditionally been based on a small number of morphological characters – primarily, the color and maculation patterns on the dorsal surface and the distribution of hairs (*setae*) on the head and body. A large number of species in the tribe Cicindelini exhibit considerable variability in color and marking (*maculation*) phenotypes. In some cases, there can be pronounced phenotypic variance within a single local population. For widely distributed species, many geographic variants have been named as subspecies (Pearson et al. 2015). A re-assessment of the species and subspecies-level taxonomy will necessarily lead to a re-evaluation of the characters traditionally used to delineate these taxa.

Morphology can be similarly misleading at the subspecies level. Subspecies in tiger beetles are typically delineated based on a limited number of characters, primarily the extent of maculations and color (usually dorsal). These characters may be under selection making them especially poor indicators of historical breaks in gene flow. Previous studies have shown that expanded white maculations can be associated with thermoregulatory abilities (Acorn 1992, Hadley *et al*. 1992) or camouflage from predators (Nayuta & Teiji 2020) and, therefore, are likely to be highly adaptive. It is expected that convergence of pattern and color could occur in separated populations due to local selection pressure of similar environmental factors. In some cases, a presumed subspecies may in fact constitute a separate species that had been wrongly placed due to the convergence of maculation and color patterns (Morgan et al. 2000, Duran et al. 2020, French et al. 2021). In other cases, variation in maculations and color may result in taxonomic oversplitting (Duran et al. 2020). This mismatch is of particular concern in tiger beetles, as subspecies is the taxonomic rank that is the focus of most conservation efforts (Knisley et al. 2014).

The North American tiger beetle species *Eunota togata* (LaFerté-Sénectère, 1841) was recently revised to update the subspecies level taxonomy based largely on the common morphological characters used for tiger beetles – maculations, dorsal color, and setal patterns – as well as inferences from historical biogeography (Acciavatti 2021). The *E. togata* species complex is halophilic, where individuals are found on open substrates that often have visible white salt crystals at the surface. All subspecies of *E. togata* have significant portions of their elytra covered by white maculations (Fig 1), with some nearly entirely maculated. As such, there exists the potential for populations to be subjected to strong selection in these environments, both for thermoregulation and for crypsis (if visual predators are present). In this thorough revision, Acciavatti (2021) recognized four subspecies: (1.) *E. togata togata*, found primarily along the Gulf Coast from Tamaulipas to Florida, with a historical disjunct population in South Carolina (known only from the early-1900s), (2.) *E. togata globicollis* (Casey, 1913), an interior subspecies ranging from eastern New Mexico and western Texas, north to Colorado and east to Oklahoma to central Kansas, (3.) *E. togata latilabris* (Willis, 1967), known only from northern Kansas and eastern Nebraska, and (4.) the newly described *E. togata leucophasma* Acciavatti, 2021, found in far western Texas, southern and central New Mexico. In addition to these four subspecies, Acciavatti (2021) reported phenotypic intergrades between *E. t. globicollis* and *E. t. leucophasma* in parts of southeastern New Mexico and western Texas.

**Figure 1.**
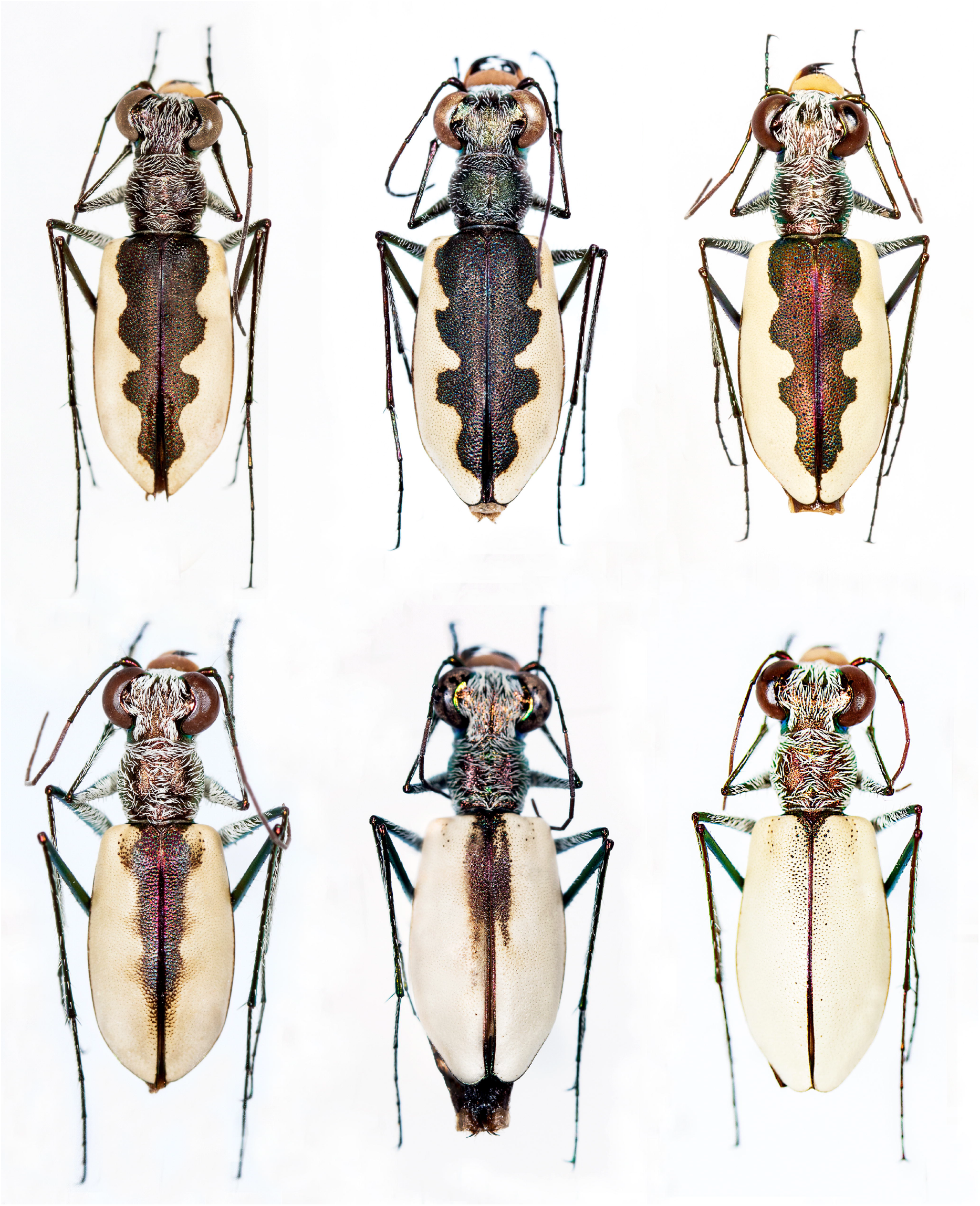
Dorsal habitus of representative populations within the *E. togata* group. The following subspecies names follow the nominal taxa recognized in Acciavatti (2021) based on morphology that are tested with genomic data in the current study. Top row (left to right): *E. t. togata* (Sour Lake, Texas), *E. t. latilabris* (Talmo, Kansas), *E. t. globicollis* (Eldorado, Oklahoma); Bottom row (left to right): *E. t. globicollis* x *E. t. leucophasma* (Laguna Gatuna, New Mexico), *E. t. leucophasma* (Laguna Del Perro, New Mexico) and *E. t, leucophasma* (Salt Flat, Texas).

In the present study, we tested these recent morphological hypotheses proposed by Acciavatti (2021) using multi-locus genomic and mtDNA analyses. We found support for some of these taxonomic hypotheses, but not others, and our phylogenetic results indicate substantial incongruence between the taxonomy and the major clades. These clades were found to correlate more strongly with geography than phenotype.

## Materials and Methods

### Specimen collection

Historical localities for *Eunota togata* were obtained from published records and publicly available online data sources, including iNaturalist, BugGuide and Symbiota Collections of Arthropods Network (SCAN). Collecting efforts were conducted to include all nominal subspecies and as many geographic areas as possible (Figure 2). All specimens and their collection localities are indicated in Supplemental Information, Table S1.

**Figure 2.**
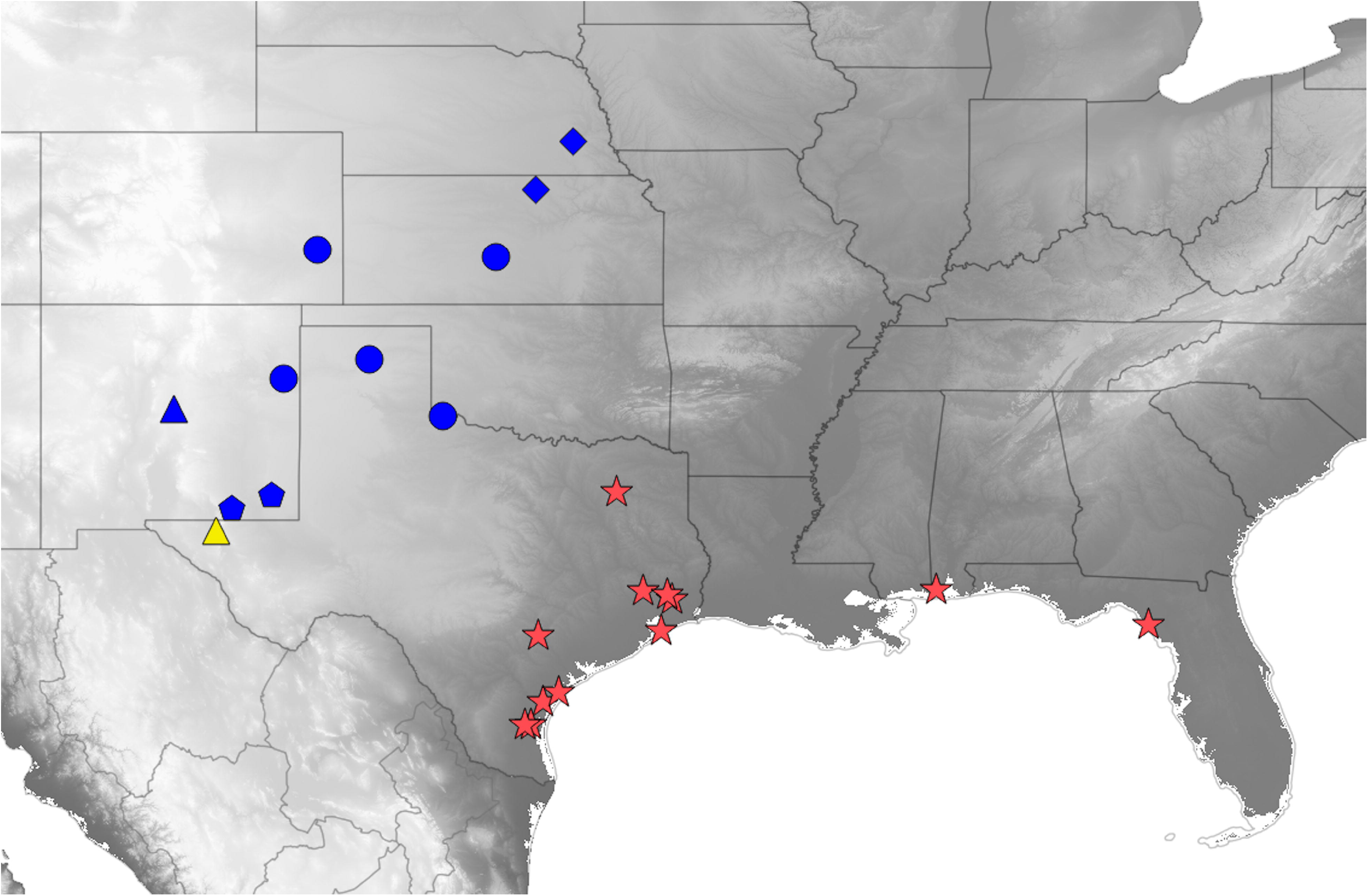
Sampling localities for the *Eunota togata* group sampled for this study. Symbols indicate existing taxonomic names for subspecies from Acciavatti (2021). Diamonds: *E. t. latilabris*, circles: *E. t. globicollis*, triangles: *E. t. leucophasma*, pentagons: *E. t. globicollis* x *E. t. leucophasma* hybrids, stars: *E. t. togata*. Colors correspond to the species supported in the present study based on mtDNA and multilocus genomic analyses. Red: *E. togata*, new status, blue: *E. globicollis*, new status, yellow: *E. leucophasma*, new status. See Table S1 for additional details of sampling localities.

### Molecular Sampling, mtDNA

Field collected specimens (preserved directly in ∼96% ethanol) and when possible, pinned specimens from private collections were sampled for molecular data. Non-destructive DNA extractions were performed in order to preserve whole specimens for morphological observations. We dried samples stored in ethanol through vacuum centrifugation until all ethanol had evaporated and then used size #3 insect pins to make 5-10 holes in the lower abdomen of each specimen. We then left intact specimens to incubate overnight at 56 ºC during the tissue lysing step of the DNeasy Blood and Tissue Kits (QIAGEN, Venlo, Netherlands) extraction protocol. Individuals were removed from lysis solution and stored in 70% ethanol before completing the DNA extraction protocol.

We used CB1 and CB2 primers (Crozier & Crozier 1992) to amplify a 424 bp region of the mitochondrial cytochrome b gene (cytb). We chose this gene due to its short fragment length, which would be more likely to amplify from degraded DNA from old or pinned specimens. The primer sequences were CB1: 5’ TAT GTW YTA CCA TGA GGA CAA ATA TC 3’ and CB2: 5’ ATW ACW CCT CCT AAT TTA TTA GGA AT 3’. PCR conditions for these primers were as follows: 2 min at 96 °C followed by 10 cycles of denaturation at 96 °C for 30 s, annealing at 46 °C for 30 s and extension at 72 °C for 1 min, then followed by 30 cycles of denaturation at 96 °C for 30 s, annealing at 48 °C for 30 s and extension at 72 °C for 1 min, with a final extension step at 72 °C for 5 min. We ran PCR reactions with a volume of 25 μL, with 12.5 μL of Apex 2X RED Taq Master Mix (Genesee Scientific, San Diego, USA), 1 μL of the forward and reverse primer (5 μM), 9.5 μL of RNase-free water, and 1 μL of DNA template. PCR product was then sent to the University of Arizona Genomics Core (Tucson, AZ) for cleanup and single-read Sanger sequencing. Sequences were deposited in the NCBI GenBank Database under the accession numbers indicated in the Appendix.

### Mitochondrial Analysis

We inferred a mitochondrial genealogy with IQ-TREE v.1.6.9 (Nguyen et al. 2015). Model selection was performed using ModelFinder in IQ-TREE with the best model chosen using BIC (Kalyaanamoorthy 2017). The tree with the best maximum likelihood score was selected from 200 independent searches. For each of the 200 runs, we estimated nodal support using 1000 ultrafast bootstraps and 1000 SH-aLRT tests. We used the -bnni command to avoid severe model violation resulting in overestimation of nodal support when performing ultrafast bootstraps. DNaSP 6.12 was used to calculate pairwise divergences between taxonomic groups. We created a haplotype network using POPART 1.7 (Leigh and Bryant 2015, Clement et al. 2002) to visualize the distribution of ancestral alleles (internal to the network) and derived alleles (tips of the network). We used the Templeton, Crandall and Sing (1992) (TCS) method for inferring a network, as this has been used extensively with mitochondrial nucleotide sequence data to estimate relationships across a wide range of genetic divergence (e.g., Gerber & Templeton 1996, Gomez-Zurita et al. 2000, Tan et al. 2020).

### Multilocus Marker Generation and Analysis

A genotype-by-sequencing (GBS) approach was used to generate multilocus nuclear markers. Specifically, we used a restriction enzyme associated DNA sequencing (RADseq) procedure to produce reduced complexity libraries as described in Parchman et al. (2012) and utilized previously for tiger beetles (Duran et al. 2019, Duran et al. 2020). RADseq libraries were sent for sequencing at the University of Texas Genomic Sequencing and Analysis Facility (Austin, TX). Sequencing was conducted in two parts. A primary sequencing run on an Illumina NovaSeq platform in a single SP lane produced 82,037,591 100 bp raw single end reads. A secondary run, on an Illumina NextSeq 500 platform, produced an additional 41,119,679 100 bp single end reads. The secondary run was conducted to include 7 additional samples in this study (TX182, TX183, TX184, AL192, AL193, AL194, and AL195) that were not part of the first sequencing run.

Ipyrad version 0.9.84 (https://ipyrad.readthedocs.io/) is a GBS toolkit for sequence assembly and analysis and was used to process all reads (Eaton 2014). Reads from each sequencing run were demultiplexed separately to reduce the number of reads assigned incorrectly. Sequences then underwent an initial filtering step where any reads with more than five base calls that had a Phred-scaled quality less than 33 were removed. From here, data was divided into four subsets based on the analyses we planned to conduct as it has been shown that identifying SNPs independently in subsets of data for downstream use can reveal patterns of genetic differentiation that might otherwise go undetected if all data is processed together (Driscoe et al. 2019). The first of these subsets was for RAxML phylogenetic analyses, which included all *E. togata* subspecies (sensu Acciavatti 2021) and sampled populations (Fig 2) – as well as three *E fulgoris* (Casey 1913) individuals and an *E circumpicta* (LaFerté-Sénectère, 1841) individual as outgroups. The next two subsets were informed by the results of the mtDNA genealogy and were as follows: 1) all nominate *E. t. togata*, 2) all E. t. *globicollis*, E. t. *latilabris*, and E. t. *leucophasma*. For each of these subsets loci were identified and loci that were present in fewer than four samples, had more than 20% SNPs, or more than 8 indels were removed to exclude poor alignments. Individuals in each subset that recovered fewer than 2,000 loci were removed from the dataset with the exception of the dataset used for RaxML as this program has been shown to be more robust to the type of missing data generated in RADseq (Stamatakis 2014). Across all subsets, individuals had data from an average of 36,051 loci.

### Multi-Locus Nuclear Trees Generated from SNP Data

A maximum likelihood tree was constructed from SNP data using RAxML 8.2.12 (Stamatakis 2014). This tree included 58 individuals, including those from every population and taxonomic group analyzed in this study. We performed N = 100 bootstrap analyses followed by 10 rapid hill-climbing maximum likelihood searches from random trees utilizing the GTRGAMMA substitution model (Yang 1994). All other parameters were left as default.

### Principal Component Analysis of SNP Data

The ipyrad analysis toolkit was used to conduct all principal component analyses. Prior to these analyses, data sets underwent additional filtering to minimize missing data: loci that were present in less than 50% of individuals in each taxonomic group or present in less than 75% of individuals overall were excluded. Remaining missing data was imputed using an algorithm that randomly sampled genotypes based on the frequency of alleles within each taxonomic group. For loci with multiple SNPs, a single SNP was randomly subsampled in order to reduce potential effects of linkage on results (Dray & Josse 2015). Analyses were each repeated 25 times and the centroid of all points from each sample was plotted.

### Bayesian Clustering Analysis Using SNP Data

We used STRUCTURE v.2.3.4 to perform unsupervised assignment of individuals to K populations using a Bayesian clustering algorithm (Pritchard et al. 2000). The model identifies population membership of individuals based on patterns of linkage and Hardy–Weinberg disequilibrium. Like for Principal Component Analyses, loci that were present in less than 50% of individuals in each taxonomic group or present in less than 75% of individuals overall were excluded from each subset of data analyzed. STRUCTURE was run with values of K ranging from K = 2 to K = 6 or K = 2 to K = 8 depending on the expected populations of the subset of data analyzed. Values of K were run with 250,000 burn-in steps and 250,000 calculation steps and replicated 10 times with default parameters. Plots of mean log probability and delta K values of each model were used to identify optimal K values (Evanno et al. 2005). Models for each value of K are presented for a more comprehensive view of the genetic variation among populations (e.g., Gilbert et al. 2012, Janes et al. 2017, Driscoe et al. 2019).

### Estimating Genetic Distance Using SNP Data

Genetic distance between populations, measured as Latter’s F_ST_ (Latter, 1972), were estimated by the hierfstat R package Version 0.5-11. Population designations and comparisons were made between the following groups based on the results of RAxML phylogeny, PCAs and STRUCTURE analyses: nominate *E. t*.*togata* and other *E. togata* subspecies; *E. t. leucophasma* from Hudspeth, TX, and other non-nominate *E. togata*; nominate *E. togata* from Florida or Alabama and nominate *E. togata* from Texas; *E. t. latilabris* from Nebraska and *E. t. globicollis* from Kansas; *E. t. leucophasma* from New Mexico and E. *t. globicollis* from Colorado.

## Results

### mtDNA analyses

The mitochondrial genealogy recovered a monophyletic *E. togata* clade, with a major split between *E. t. togata* and the remaining nominal taxa (Fig. 3). Herein, we will refer to these clades as the Gulf Clade and the Interior Clade, respectively. No clear geographic or taxonomic structure existed within the Gulf Clade (i.e., *E. t. togata*). Within the Interior Clade, the first bifurcation was between the population of *E. t. leucophasma* from Salt Flat (Hudspeth County), Texas, and a clade containing all other Interior Clade populations (*E. t. globicollis, E. t. latilabris, E. leucophasma* from Laguna Del Perro [Torrance County], New Mexico, and the putative hybrids between *E. t. leucophasma* and *E. t. globicollis*). The average pairwise sequence divergence between the Gulf and Interior Clades was 5.2%. Within the Interior Clade, the divergence between the Salt Flat, Texas population of *E. t. leucophasma* (type locality) was 2.0% from the clade containing the remaining taxa.

**Figure 3.**
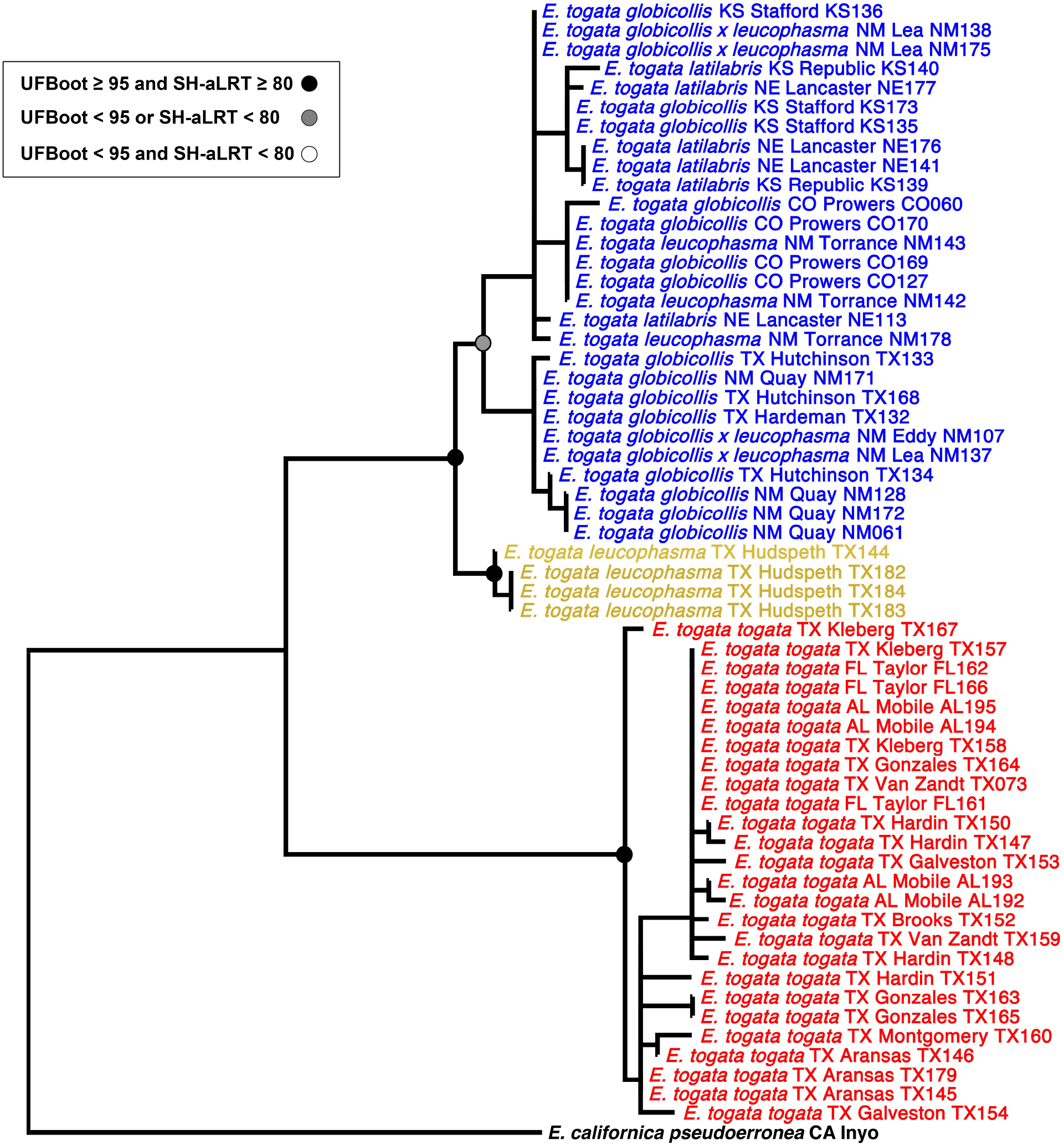
Maximum-likelihood mtDNA genealogy inferred in IQ-TREE based on cytb gene. Taxon naming follows previous naming conventions (Acciavatti 2021) with colors highlighting new taxonomic groupings that we identified in this study, colored as in Figure 2. Red: *E. togata*, new status, blue: *E. globicollis*, new status, yellow: *E. leucophasma*, new status.

The TCS mtDNA haplotype network (Fig 4) showed that the Gulf and Interior Clades were separated from each other by 14 mutational steps, the largest subdivision within the network. Within the Gulf Clade, internal (ancestral) alleles were widely geographically distributed and tip (derived) alleles were similarly distributed across the range. Within the Interior Clade, the population of *E. t. leucophasma* from Salt Flat, Texas contained the most ancestral alleles and were separated by 7-10 mutational steps from the remaining Interior Clade populations. The other major clade within the Interior Clade had ancestral alleles that were primarily distributed in the southernmost parts of the clade’s distribution - New Mexico and parts of the Texas panhandle. Derived alleles were mainly distributed in the northern part of the range: Colorado, Kansas and Nebraska.

**Figure 4.**
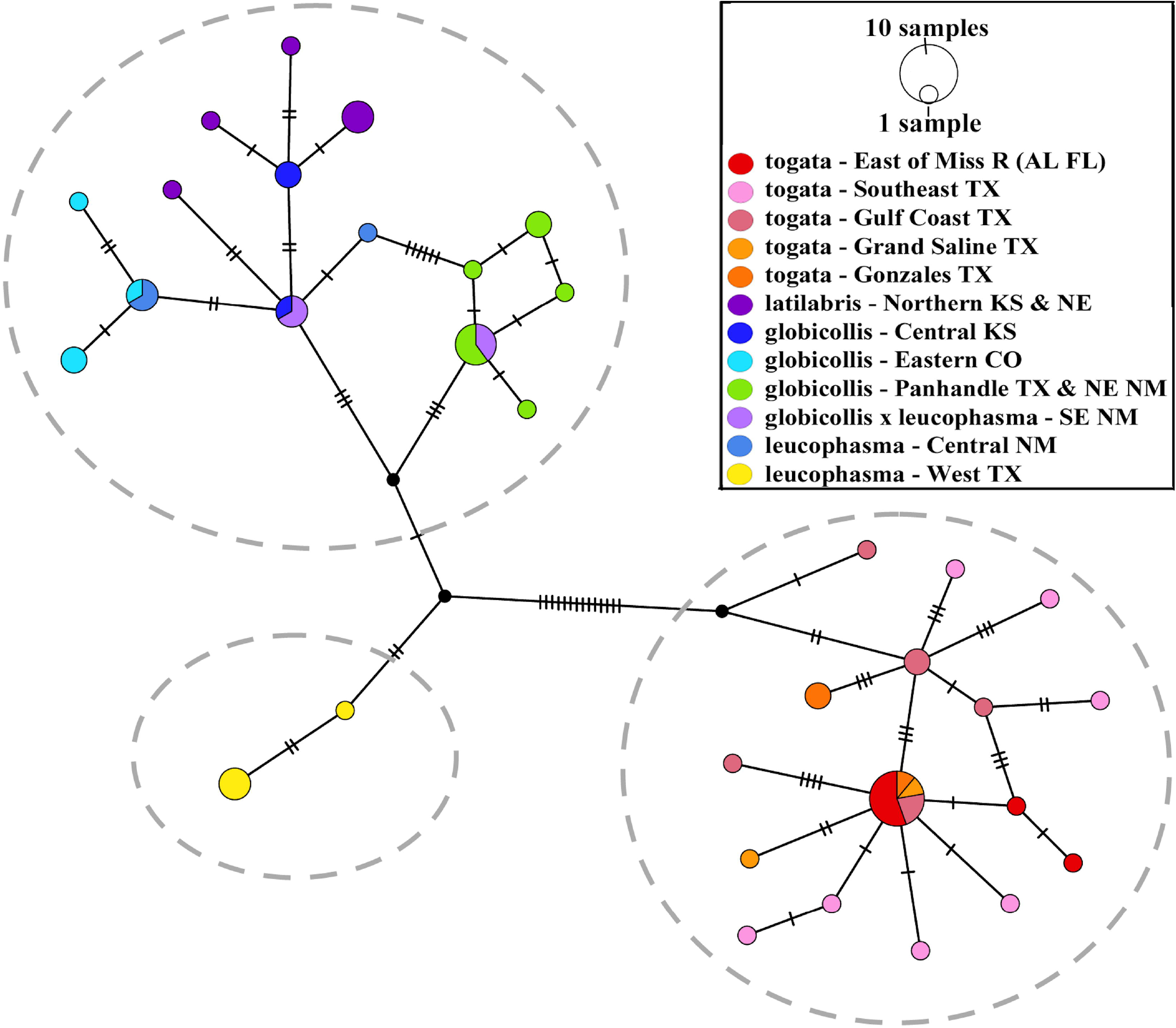
TCS haplotype network generated with POPART 1.7 (Leigh and Bryant 2015) shows the evolutionary relationships of the cytb sequences in this study. Each hatch mark indicates a mutational step between adjacent alleles. The color of each circle corresponds to the geographic location of the sequence (see figure legend). The size of the circle is proportional to the haplotype frequency.

### Maximum Likelihood Phylogeny from Genome-wide SNP Data

The RAxML tree based on 36,051 total loci recovered clades that were mostly congruent with the mtDNA genealogy (Fig. 5). The tree exhibited a major split between the Gulf and Interior Clades (Latter’s F_ST_=0.229), and another deep split within the Interior Clade that separated the Salt Flat Texas *E. t. leucophasma* and the Interior Clade containing all other populations (Latter’s F_st_=0.220). A topological difference between the RAxML tree and the mtDNA gene tree is the presence of a monophyletic group comprised of the *E. t. togata* individuals found east of the Mississippi River nested within the Gulf Clade in the RAxML tree. Moreover, this clade had a longer branch length than any other taxa within the Gulf Clade. All of the aforementioned clades were supported by bootstrap values above 95.

**Figure 5.**
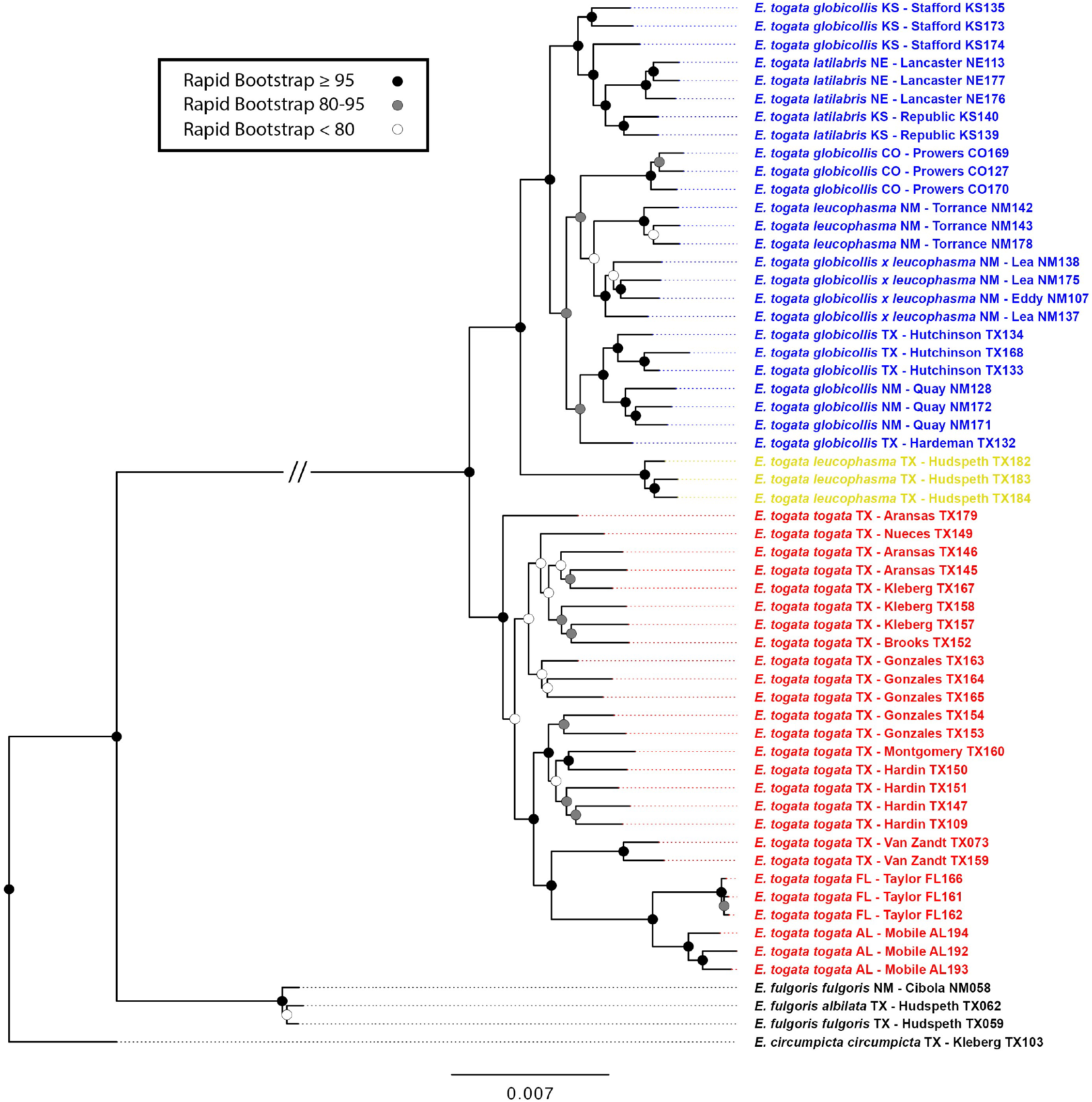
Maximum likelihood (RAxML) tree based on SNP dataset. Topology based on 36,051 total loci for the same individuals sampled in the mtDNA genealogy. Taxon naming follows previous naming conventions (Acciavatti 2021) with colors highlighting new taxonomic groupings that we identified in this study, colored as in Figure 2,3. Red: *E. togata*, blue: *E. globicollis*, new combination, yellow: *E. leucophasma*, new combination. The severed branch indicated by “//” represents a shortened branch length of 0.028.

### PCA from Genome-wide SNP Data

A principal component analysis of the SNP data was conducted to assess the clustering of individuals within each of the two deeply separated clades, the Gulf Clade (i.e. *E. t. togata*) and the Interior Clade (i.e. all other subspecies and putative hybrids) (Figure 6A,B). For the Gulf Clade, the first two principal components explained 27.1% of the total variation in the dataset. The observed results indicated a substantial separation between the individuals east and west of the Mississippi River. For the Interior Clade, the first two principal components explained 26.2% of the total variation in the dataset. The largest separation was between the Salt Flat, Texas *E. t. leucophasma* and all other individuals, including the New Mexico *E. t. leucophasma*. Other populations were spread along the PC1 axis consistent with a latitudinal cline, ranging from New Mexico *E. t. globicollis* (bottom of PC1) to Nebraska *E. t. latilabris* (top of PC1).

**Figure 6.**
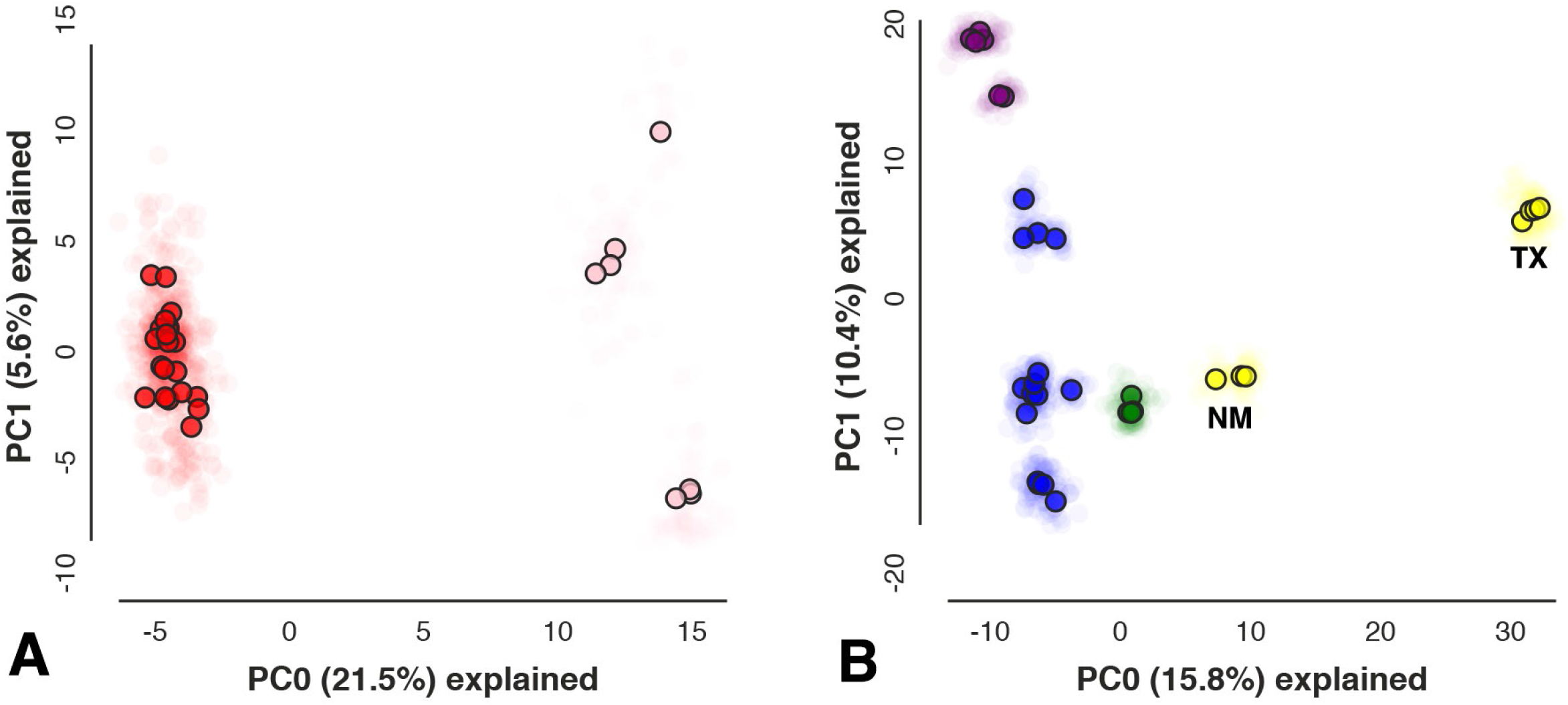
Principal component analyses (PCA) of 2,269 (A) and 5,736 (B) SNPs. Loci were limited to those found in a minimum of 50% of individuals in each nominal taxonomic group and in 75% of individuals overall to produce a SNP matrix with relatively little missing data (19.08%; 16.85% for A and B respectively). Transparent points represent replicate analyses (N = 25) while opaque points represent the centroids of these replicates. Panel (**A):** PCA of Gulf Clade individuals (*E. t. togata*). Red points on left-hand side of graph corresponds to *E. t. togata* individuals from west of the Mississippi River. Pink points on the right correspond to individuals from East of the river. Panel (**B):** PCA of Interior Clade individuals (all remaining *E. togata* subspecies). Blue points correspond to *E. t. globicollis*, purple points correspond to *E. t. latilabris*, yellow points correspond to *E. t. leucophasma*, and green points correspond to phenotypic intergrades between *E. t. globicollis* and *E. t. leucophasma*. New Mexico *E. t. leucophasma* are recovered closer to other populations of the Interior Clade than to the Texas *E. t. leucophasma*. Populations on the left side of the graph exhibit a latitudinal cline from New Mexico *E. t. globicollis* (bottom) to *E. t. latilabris* from Nebraska (top).

### Bayesian Clustering Analysis Using Genome-wide SNP Data

Based on the results of the multilocus phylogeny, PCAs, and the mtDNA genealogies, we conducted a set of Bayesian clustering analyses using the program STRUCTURE v.2.3.4 to assess the genetic structuring within the Gulf and Interior Clades (Fig. 7). Within the Gulf Clade, at K=2, the two groups identified corresponded to the populations of *E. t. togata* west and east of the Mississippi River. At all additional K populations, the identified groups did not change significantly. Within the Interior Clade, at K=2, the two groups identified correspond to the Salt Flat Texas *E. t. leucophasma* and the remaining Interior Clade. At K=3, an additional group was identified that corresponded to *E. t. latilabris* populations and the central Kansas *E. t. globicollis*. At K=4, there is a partial separation of the *E. t. globicollis* individuals from Colorado. At all additional K populations, the identified groups did not change significantly.

**Figure 7.**
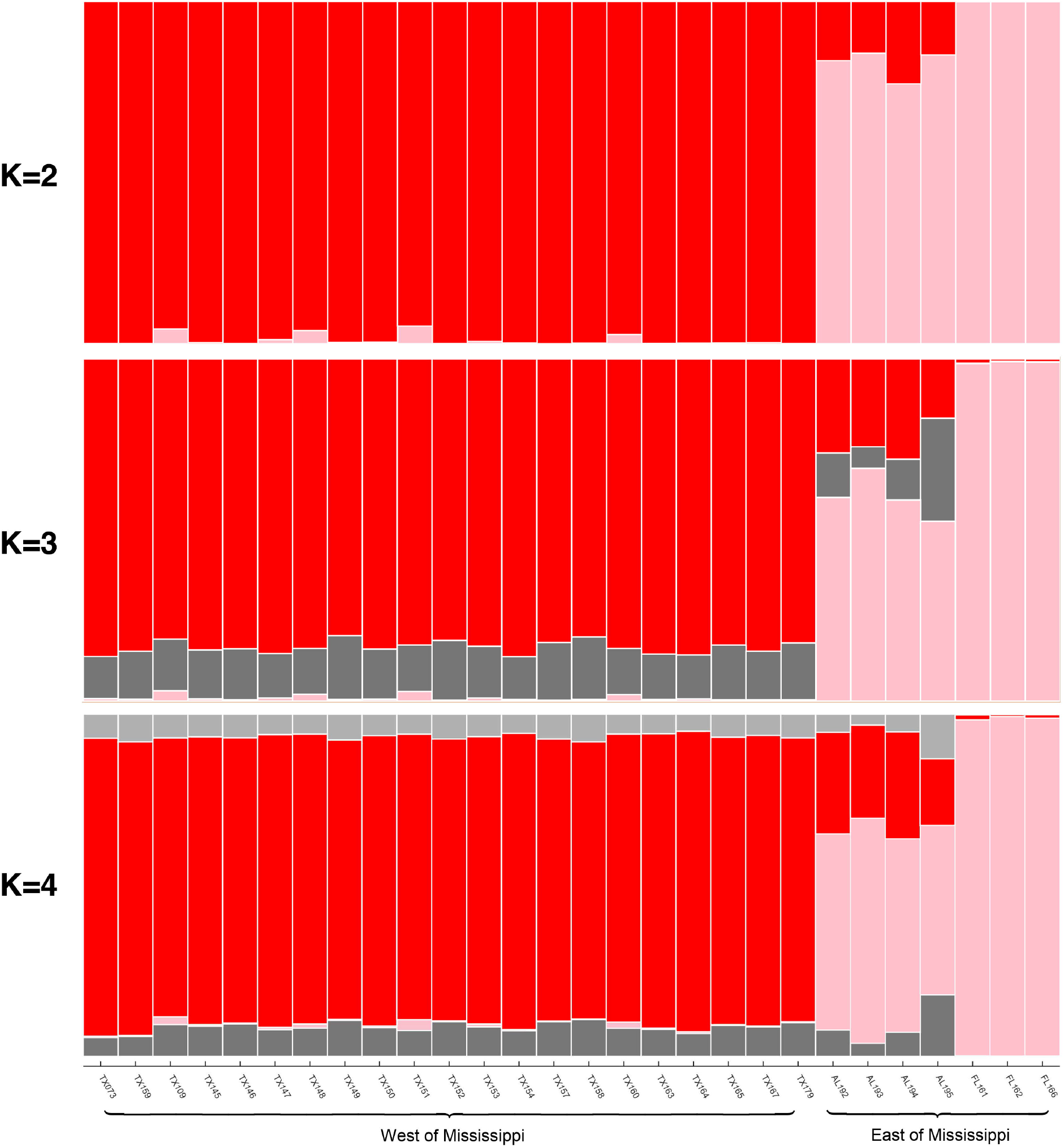
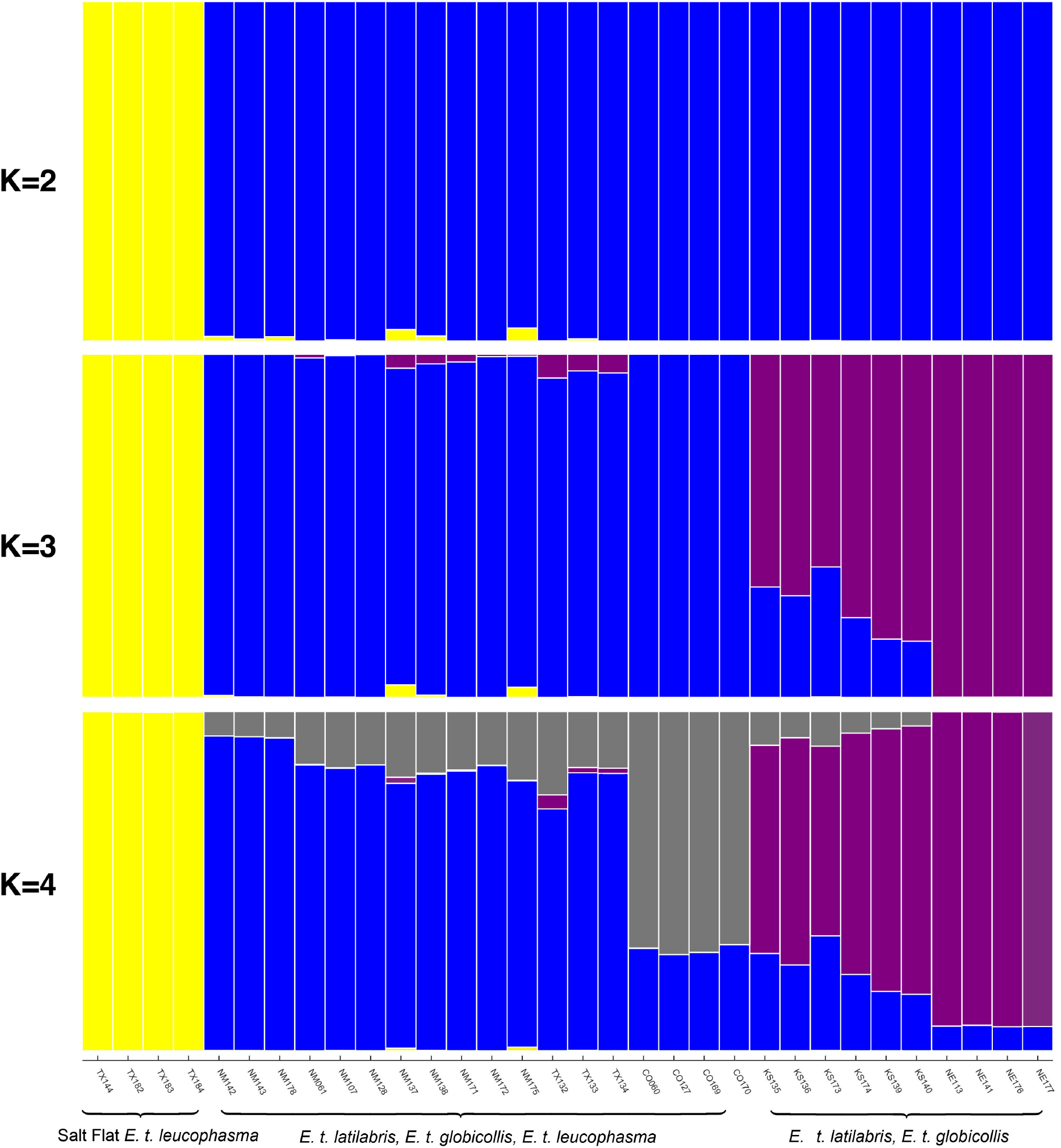
STRUCTURE analyses of 2,269 (A) and 5,915 (B) SNPs. Loci were limited to those found in a minimum of 50% of individuals in each taxonomic group and in 75% of individuals overall to produce a SNP matrix with relatively little missing data (19.08%; 17.02% for A and B respectively). Delta-K and mean log probability plots are illustrated in Figure S1. A) K=2-4 for Gulf Clade (i.e. *E. t. togata*), B) K=2-4 for Interior Clade (all other subspecies and intergrades)

### Taxonomy

Only one of the four taxonomic entities recognized in Acciavatti (2021) is fully supported by our analyses. *Eunota togata togata* was always recovered as a distinct clade (Figs. 3-5) consistent with the way it was circumscribed. The taxonomic entities referred to as *E. t. globicollis* and *E. t. leucophasma* fall into two distinct clades (Figs 3-7). Both of these groupings are different from those circumscribed in the previous revision. We found that the primarily *E. t. globicollis* clade included populations from within the previously recognized range (eastern New Mexico and Colorado to central Kansas), but also included “intergrade” or “*E. t. globicollis* x *E. t. leucophasma*” populations from eastern New Mexico.

Additionally, the *E. t. globicollis* clade contained the populations from northern Kansas and eastern Nebraska that were known as *E. t. latilabris* in Acciavatti (2021). As such, we recognize *E. t. latilabris* as a new synonym. The distinct *E. t. leucophasma* clade consisted of only one population, those from the type locality of Salt Flat, Texas. The other population that Acciavatti (2021) recognized as *E. t. leucophasma*, from Laguna Del Perro, New Mexico, was contained within the *E. t. globicollis* clade.

Given the plurality of results, including reciprocal monophyly and large genetic divergences between the three major clades, we recognize *E. t. togata* as *E. togata* and elevate *E. t. globicollis* to a full species, *E. globicollis* and *E. t. leucophasma* from Salt Flat, Texas, to a full species, *E. leucophasma*. Additionally, *E. togata* populations east and west of the Mississippi River were observed to be partially differentiated based on the PCA (Fig 6) and STRUCTURE analyses (Fig 7), but not in the mtDNA genealogy (Fig 3) and were found to be a distinct clade, but not reciprocally monophyletic in the RAxML multilocus tree (Fig 5). As such, the eastern populations of *E. togata* may be considered a new subspecies, but this will require further study.

Given our results, three distinct species are formally recognized in the *Eunota togata* group. They are as follows:

***Eunota togata* (LaFerté-Sénectère 1841), stat. nov**.

***Eunota globicollis* (Casey 1913), stat. nov**.

***Eunota leucophasma* (Acciavatti 2021), stat. nov**.

And we synonymize the following taxon:

***Eunota latilabris* (Willis 1967), syn. nov**.

## Discussion

### Discordance between genetic analyses and morphological taxonomy

Our results indicate that there is considerable discordance between the morphologically defined subspecies (Fig 1) and the clades we identified with multiple genetic methods based on mtDNA and multilocus genomic data. Color and maculation patterns were primarily used to circumscribe the subspecific taxa within *E. togata* (Acciavatti 2021), including the heavily maculated populations referred to as *E. t. leucophasma* that were found in west Texas, south central New Mexico, and in a disjunct set of populations in central New Mexico. Despite morphological similarity (both sets of populations have nearly entirely white elytra), we demonstrate that they are distantly related. In the mtDNA genealogy, TCS haplotype network, RAxML multilocus tree, PCA, and STRUCTURE analyses, they were always found in separate clades or groups. The degree of genetic differentiation between these two nominally *E. t. leucophasma* populations was high (Latter’s F_ST_=0.220) whereas by comparison, the differentiation between New Mexico *E. t. leucophasma* and Colorado *E. t. globicollis* was much lower (Latter’s F_ST_=0.037). This demonstrates that the nearly all white maculation pattern has evolved independently at least two times in this group.

*Eunota t. latilabris* (defined as populations from Nebraska and northern Kansas Great Plains) was believed to be distinct from *E. t. globicollis* (central, southern Kansas, Colorado, Oklahoma, Texas Great Plains) largely based on coloration, and a potential explanation for this evolutionary division was given. Acciavatti (2021) postulated that the *E. t. latilabris* populations were dark olive green to dark brown because they evolved in small areas of dark saline soil habitats in northern parts of the range of *E. togata*, contrasting with redder Permian soil deposits to the south that correspond with the more coppery-red dorsal colors of *E. t. globicollis*. Our results do not support the hypothesis that color correlates with subdivisions of the species in the Great Plains region. *Eunota t. latilabris* are not monophyletic in either the mtDNA genealogy or the multilocus phylogeny. Individuals of *E. t. latilabris* group together in the PCA of SNPs, but this is also consistent with geography and isolation-by-distance, as the *E. t. latilabris* and *E. t. globicollis* populations form a North-South cline alone the PC1 axis (Fig. 6). Additionally, a subdivision was observed in the multilocus phylogeny and the STRUCTURE analyses that included Nebraska and northern Kansas *E. t. latilabris* as well as central Kansas *E. t. globicollis*. This genetic group spanned across the taxonomic boundary and was not consistent with the recent revision based on coloration.

Some populations display maculation patterns that appear to be intermediate between *E. t. globicollis* and the more heavily maculated *E. t. leucophasma* and these had been treated as intergrades between the two otherwise phenotypically distinct subspecies (Acciavatti 2021). Our analyses demonstrates that these partially expanded maculations were poor predictors of relatedness, as these populations were always found nested within clades or clusters of *E. t. globicollis*. No support for hybridization or intergradation was observed with STRUCTURE. If these phenotypically intermediate populations were the result of intergradation between *E. t. globicollis* and *E. t. leucophasma*, this would have been apparent in the STRUCTURE results, where a pattern of admixture in these individuals should have been observed.

A potential limitation of any population-level phylogenetic study is sampling. Although we sampled all nominal taxa (sensu Acciavatti 2021) and had broad geographic coverage across the *E. togata* group range, it is possible that addition samples would yield additional inferences. For example, we did not have material from White Sands, NM, although a population of the *E. togata* group has been recorded from that area. Given the historical biogeography (discussed below), it is very likely that this population will belong to *E. leucophasma*, stat. nov. Still, additional sampling would not change any of the major inferences, but could lead to additional ones.

### Discordance between mtDNA and genomic analyses

There was broad agreement between the results of the mtDNA analyses and the multilocus genomic analyses. Each of the distinct species that we inferred via this study, *E. togata*, stat. nov., *E. globicollis*, stat. nov. and *E. leucophasma*, stat. nov. were supported by all mtDNA and multilocus analyses. However, more genetic structuring was observed in the multilocus tree than in the mtDNA genealogy. For example, a long-branch clade that included all of the *E. togata* stat. nov. east of the Mississippi was present in the multilocus phylogeny (Fig 5), but absent in the mtDNA tree (Fig 3). Similarly, a clade containing all *E. globicollis* stat. nov. populations from central and northern Kansas and eastern Nebraska was recovered in the multilocus phylogeny, but not in the mtDNA tree.

Inferences based on mtDNA alone can be contentious, as the phylogenetic patterns based on this marker can be inconsistent with those derived from the rest of the genome (Funk & Omland 2003, Ballard & Whitlock 2004, Gompert et al. 2006, Toews & Brelsford 2012, Filée et al. 2022). In some cases, mtDNA may fail to accurately delineate closely related species, especially when hybridization and introgression occur (Fitzpatrick et al. 2010, Duran et al. 2020). Although the mtDNA analyses did recover the same species as the multilocus ones, it is curious that there was reduced structure in the mitochondrial topology. One possible explanation is the purging of genetic variation due to purifying natural selection on mtDNA (reviewed in Sun et al. 2018). An alternative explanation is the role of the maternally-inherited, reproductive parasite, *Wolbachia*, a gram-negative endosymbiotic bacteria that can manipulate reproduction, sex ratios, and inheritance (Werren et al. 1995). Future studies of the Gulf Coast populations of *E. togata* may elucidate the nature of this mitochondrial similarity east and west of the Mississippi River generating this mito-nuclear discordance.

### Historical biogeography of the entire E. togata species complex

The three major lineages recovered in this study each correspond to a different geographic area (Fig 2), presumably due to allopatric speciation in these regions. *Eunota togata*, stat. nov. (=Gulf Clade) is considerably separated from the Interior Clade containing the other two species, *E. globicollis*, stat. nov. and *E. leucophasma*, stat. nov. The large degree of genetic differentiation (mtDNA pairwise divergence = 5.2%; Latter’s F_ST_ from multilocus genomic data = 0.229) separating *E. togata*, stat. nov. from the others underscores that this separation of ancestral *togata* group populations between low elevation, primarily coastal salt flats and more elevated interior saline and alkaline habitats must have been an ancient event. Given this genetic divergence, if we apply an approximate molecular clock assuming mtDNA genes diverge about 2.3% per million years (i.e., Brower 1994), the divergence between the Gulf and Interior Clades would likely have happened in the late Pliocene to early Pleistocene. Linking these newly recognized lineages with biogeographic patterns more locally may lead to addition inferences on the evolution within each group. Most notably, there are two major biogeographic anomalies that stand out in the results of this current study: (1.) inland populations of *E. togata, stat. nov*., on salt domes appear to be disjunct from the coastal populations by up to 300 km, and (2.) *E. leucophasma, stat. nov*., as defined here is endemic to the Salt Flat basins near the Guadalupe Mountains of West Texas and distinct from the other light-colored morphs in *E. globicollis*, stat. nov. from the High Plains.

#### (a.) Biogeography of the Coastal Clade – E. togata, stat. nov

*E. togata*, stat. nov. is confined primarily to the coastal plain along the Gulf Coast. Along the immediate coast, localized alkaline playas develop on the landward side of barrier islands. These barrier islands are largely associated with relative sea level transgression since the last glacial maximum approximately 20,000 years ago (Milliken et al. 2008, Anderson et al. 2022). The back bay behind the barrier islands are highly dynamic environments, oscillating between estuarine to full isolation from the ocean. Playas also exist 10-100 km inland of the immediate coast in East Texas. These are not directly related to current coastal processes but are associated with local subsidence, leading to the formation of small saline playas.

Further inland populations of *E. togata* on salt domes, such as Grand Saline, TX (Figure 2; Table S1), appear to be disjunct from the coastal populations by up to 300 km. These inland populations are associated with saline prairies that replicate the conditions of coastal saline flats because of the influence of salt domes. However, if populations of *E. togata* have been reproductively isolated for two million years on salt domes then we would expect to see genetic differences equivalent to those observed in the far West Texas populations of *E. leucophasma*. The lack of such divergence indicates that inland populations of *E. togata* probably result from more recent, constant dispersal events. It may be possible that disjunct populations of *E. togata* are the result of long-distance dispersal over unfavorable habitat or that there were times when saline habitats extended down coastal rivers to connect with the coast.

Presently the Brazos, Red and Pecos Rivers exhibit high levels of saline water. The Sabine River is one potential corridor for tiger beetle dispersal historically. It presently communicates with the intermittent salt sources of Jurassic domes, but not the Permian salts of Oklahoma, New Mexico and West Texas. But there is evidence that sometime since the Miocene, the Sabine did make this connection. Salt inflow to the Sabine has been interrupted by the pirating of the Red River, which has at some point captured the upper Sabine (Manning 1990).

#### (b.) Biogeography of the Interior Clade – E. leucophasma, stat. nov

*Eunota leucophasma*, stat. nov, from Salt Flat (Hudspeth Co), Texas, appear to have diverged from the rest of the Interior Clade early in the evolution of this group (mtDNA pairwise divergence = 2.0%; Latter’s F_ST_ from multilocus data = 0.220). This may be due to the unique geology of the Salt Flat Playa, which is about 240 km long and 8 to 24 km wide, making it one of the largest gypsum playas in the United States. *E. leucophasma*, stat. nov, occurs in the Salt and Tularosa Basins in west Texas and southern New Mexico, respectively. Both of these basins are part of the Rio Grande watershed, defined tectonically by the Rio Grande Rift. The Rio Grande Rift is a north-south trending depression extending from northern New Mexico-southern Colorado south through south-central New Mexico and west Texas and north-central Mexico. This depression is generated by continental rifting that initiated ∼25 million years ago, continuing to the present. Such rifting produced a series of rift basins, ranging from the Santa Fe basin in north-central New Mexico to the Tularosa Basin and Salt Basin in the southern Rio Grande rift. Extension accelerated between 3 to 8 million years ago (Kelley 1979, Seager and Morgan 1979, Abbey and Niemi 2020). The Rio Grande Rift is far enough east from the Pacific that it lies in the rain shadow of the Sierra Nevada and Peninsular Ranges in California, and far enough west from the Gulf of Mexico that moisture-laden air from the Gulf does not reach the region except during late summer monsoons. The land-locked nature of the basins, coupled with low precipitation, result in the ideal conditions for generating alkaline playas. These rift basins are isolated from the High Plains, where *E. globicollis* resides, by a topographic high. The basins in southern New Mexico and west Texas are flanked by mountains composed of Paleozoic marine sediments deposited in inland sea-like environments (e.g., the Delaware Basin of the Guadalupe Mountains; Hills 1984). Those ancient depositional environments were conducive to forming evaporite gypsum deposits. Erosion and weathering of these Paleozoic gypsum deposits provide a ready source of gypsum into these basins, which are reworked by wind in the arid conditions, resulting in white gypsum playas and sand dunes (e.g., White Sands National Park in the Tularosa Basin; Mamer and Newton 2017). Such gypsiferous soils are not found in the ranges of *E. globicollis* or *E. togata*. In summary, despite the proximity of *E. leucophasma* in the Salt Basin to occurrences of *E. globicollis* in the High Plains, these rift basins appear to have been separated from the High Plains by a long-lived (My) topographic high.

In extreme environments, such as the Salt Flat Playa, the organisms that live there may be highly specialized (Powell 1980 Hussain 1998). Pupfishes (genus *Cyprindon*) live in isolated desert waters ranging across the Basin and Range Province from California to western Texas. Their diversification is thought to have been driven by the drying of Pleistocene lakes and streams as extreme chemical and physical conditions together with allopatry led to the evolution of tolerances of salinity three to four times that of sea water and temperatures up to 45 ºC (113 ºF; Miller 1981). Rodents that inhabit the salt flat basins in Argentina have been shown to possess adaptive traits more similar to the kangaroo rats (*Dipodomys sp*.) that inhabit the sand dunes near Salt Flat, TX than more closely related members of their own infraorder (Hystricognathi; Ojeda et al. 1999, Guadalupe Mountains National Park Survey of Sand Dune Mammals). For *E. leucophasma*, stat. nov., the extreme habitat in the Salt Flat Playa may have played an important role in their divergence from the common ancestor shared with *E. globicollis*, stat. nov., most notably the evolution of their distinctive white maculations, which cover the entirety of their elytra and thus may provide camouflage from predators. This would be concordant with similar patterns of color evolution seen in animals endemic to habitats with uniquely colored sediments, such as lizard species on the gypsum dunes of White Sands National Monument, New Mexico (Rosenblum 2006) or light colored beach mice on the coast of Florida (Mullen et al. 2009, Vignieri et al. 2010). Moreover, there is strong evidence that white maculations have thermoregulatory effects (Acorn 1992, Hadley et al. 1992) and as such, the *E. leucophasma*, stat. nov. on the expansive white Salt Flat Playa may benefit from expanded unpigmented areas. This may also explain how the populations of *E. globicollis* stat. nov. from Laguna del Perro in central New Mexico (previously treated as *E. t. leucophasma*; Acciavatti 2021) may have converged on expanded white maculations as they also live on one of the largest white playas in the southwestern US.

#### (c.) Biogeography of the interior clade – E. globicollis stat. nov

*Eunota globicollis*, stat. nov. appears to be restricted to the High Plains physiographic province. In New Mexico, even though this clade occurs within 100 km of *E. leucophasma*, stat. nov., *E. globicollis*, stat. nov. occurs strictly east of the continental divide in the High Plains while *E. leucophasma*, stat. nov. is restricted to the Rio Grande Rift (Figure 2). The High Plains form a gradual slope flanking the east side of the Rocky Mountains and extend from west Texas north to South Dakota. A unique feature of the High Plains is its abundance of small playas, which form some of the most important wetlands for migratory birds in North America. From 20 to 5 million years ago, uplift of the Rocky Mountains to the east led to a large sloping fan of sediments that make up the substrate of the High Plains (Willet et al. 2018). Uplift, however, appears to have progressively swept eastward such that regions that were once depositional transitioned into an erosional phase, resulting in a dynamic transient geomorphology (Willet et al. 2018). Because of the transient nature of the geomorphic systems, the High Plains is characterized by regions of incision as well as regions of relative stasis that have not yet been influenced by incision. These stable regions develop calcrete layers in the soil, which later undergo dissolution, resulting in localized depressions, not unlike karst systems.

Of interest here is lighter morph *E. globicollis*, stat. nov. existing at Laguna del Perro in the Estancia Basin, which was previously described as *E. t. leucophasma*. However, this divergence is consistent with the geological history of this region as the Estancia Basin lies to the east and outside of the Rio Grande rift basin and is physiographically more similar to the High Plains. The Estancia Basin is a relict of a Paleozoic basin, but now sits as a shallow, internally drained basin at 1.8-3 km in elevation. The Estancia Basin is separated by the High Plains by weak topographic relief, but separated from the Rio Grande rift by a north-south trending series of rugged mountains on the east flank of the Rio Grande rift, thus, migration between the Estancia Basin and the Rio Grande rift basins should have been unlikely.

## Conclusions

Our study highlights the importance of reevaluating the morphology-based taxonomy of conservationally important species using multiple complementary datasets to more precisely characterize the units of biodiversity. Phenotypic variation corresponding to classic morphological subspecies delineations was not closely associated with patterns of genetic variation. We found that several color-based subspecies were not supported as distinct evolutionary units. In contrast, our analyses indicated that the species-level diversity was underestimated. Previously, *Eunota togata* was a single species with four recognized subspecies (Acciavatti 2021). Our analyses indicated that there were three distinct species within this taxonomic group. These results demonstrate the need to critically re-evaluate the role of phenotypic inference for the taxonomy of this popular group of insects.

## Acknowledgements

The authors express their gratitude for the generous support of those who made this study possible. For assistance in collecting of specimens, we thank Stephen Spomer, Harlan Gough and Aaron Chambers. We thank Zach Holzman for help with Figures 4 and 7. We thank two anonymous reviewers for their constructive input on the manuscript.

